# Porosity-permeability tensor relationship of closely and randomly packed fibrous biomaterials and biological tissues: Application to the brain white matter

**DOI:** 10.1101/2023.01.14.524065

**Authors:** Tian Yuan, Li Shen, Daniele Dini

## Abstract

The constitutive model for the porosity-permeability relationship is a powerful tool to estimate and design the transport properties of porous materials, and has thus attracted a significant number of attention for the advancement of composite materials. However, in comparison with engineering composite materials, biomaterials, especially natural and artificial tissues, have more complex micro-structures such as high anisotropy, high randomness of cell/fibre dimensions and very low porosity. Consequently, these properties of the biomaterial make the porosity-permeability relationship more difficult to obtain than traditional composites. To fill this gap, we start the mathematical derivation from the fundamental brain white matter (WM) formed by nerve fibres. This is augmented by a numerical characterisation and experimental validations to obtain an anisotropic permeability tensor of the brain WM as a function of the tissue porosity. Moreover, we propose an anisotropic poroelastic model enhanced by the newly defined porosity-permeability tensor relationship which precisely captures the tissue’s macro-scale permeability changes due to the micro-structural deformation in an infusion scenario. The constitutive model, the theories and protocols established in this study will both provide improved design strategies to tailor the transport properties of fibrous biomaterials and enable the non-invasive characterisation of the transport properties of biological tissues. This will lead to the provision of better patient-specific medical treatments, such as drug delivery.

## 1. Introduction

Fluid transport in porous media is a ubiquitous phenomenon in nature, which has attracted widespread attention and extensive research in the fields of geoscience & petroleum engineering [1], environmental engineering [2], composite material [3, 4], biological & medical science [5, 6] etc. The development of porous transport theories has also tremendously advanced our understanding and capability of designing biomaterials, such as the bone-substituting and -repairing biomaterials [7, 8], artificial bio-membrane [9, 10], and drug carriers [11, 12]. However, the microstructural complexity, especially in natural and artificial tissues, remains an obstacle in understanding the materials’ transport properties [13]. This is because hydraulic permeability, the key parameter to determine the transport efficiency of a porous medium, is highly dependent on the pore-level structure of the said medium. This not only prevents us from achieving high efficiency and customisation of biomaterials by direct design of their microstructure, but also hinders our capability to achieve a better understanding of the functions and properties of biological tissues.

Linking the material’s geometric properties and its permeability is regarded as a cornerstone for the improved design of porous materials [4, 14, 15]. The classical and well-known semi-empirical Kozeny-Carman (KC) formula that defines the permeability (*κ*) of saturated sands as a function of its porosity (i.e. void fraction, *ϕ*, Eq. 1) [16, 17] and its very broad application is a good example of this:

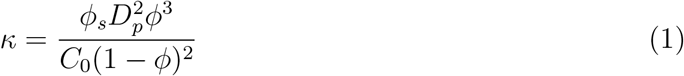

where *ϕ_s_, D_p_, ϕ* are the particle sphericity, diameter, and sample porosity, respectively; *C*_0_ is an empirical coefficient depending on the constituent geometries. With many further improvements based on theoretical and experimental techniques, the porosity-permeability relationship in isotropic porous medium has been thoroughly investigated with different formulas developed to provide satisfactory predictions in different scenarios, as reported in e.g. Refs. [18, 19].

However, microstructural anisotropy widely exists in biological tissues and biomaterials, e.g., brain, muscles, skin, cartilage, bio-gels, scaffolds [20, 21]. This adds more complexity to link microstructural properties to the macroscale permeability, as permeability can no longer be described by a scalar but becomes a tensor and all its components need to be appropriately determined. Studies on fibrous materials have put forward some formulas to relate the porosity and fibre dimensions to the permeability, such as the relationships proposed by Gebart[22] and further improved by Clague [23] and Nabovati [4], as shown in Eq. 2; alternative expressions have also been proposed by Van Doormaal [24] in Eq.3 based on the KC formula.

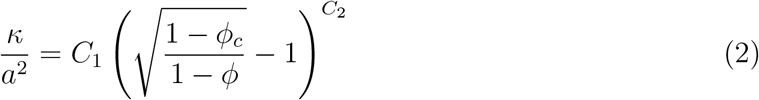

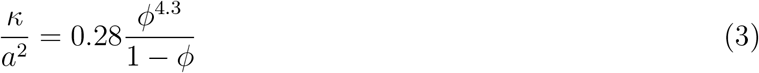

where *a* is the fibre radius, *ϕ_c_* is the critical value of porosity below which no permeating flow can develop and *C*_1_ *C*_2_ are related to the network geometry. Shou also proposed a more sophisticated form of the permeability tensor[25]. However, significant gaps still exist in applying these formulas to biomaterials, because (i) these formulas exclude strong anisotropy as fibres in composite materials often have random orientation whereas cells in some biological tissues have more uniform directionality, e.g. nerve fibres and muscle cells; (ii) these formulas were derived by adopting fibres with constant diameter, but cells and fibres in biomaterials span across a range of sizes; (iii) almost all these formulas are invalid when the porosity is lower than 0.3; however, cells and biological fibres are normally closely packaged, leading to low porosities; the porosity of brain tissue, for example, is within the 0.18 ~ 0.3 range. In addition, the limited availability of tissues makes it impractical to collect sufficient tissue samples with different porosities to validate the porosity-permeability tensor relationships. Consequently, putting forward a porosity-dependent permeability tensor suitable for fibrous biomaterials and biological tissues is challenging and has, at least until recently, eluded scientists.

In this work, we establish a porosity-permeability tensor relationship (***κ*** = *f* (*ϕ*)) of closely and randomly packed fibrous biomaterials, starting from the specific application to the brain white matter (WM). This was achieved through the integration of mathematical derivation and experimental characterisation of the microstructure of WM reconstructed from high-resolution imaging data. Furthermore, based on the newly developed porosity-permeability tensor relationship, we propose an anisotropic poroelastic model for fibrous biomaterials and validate it using ovine brain infusion experiments. In this study, we (i) develop software capable of generating the microstructure of closely and randomly packed fibrous biomaterials for microstructure-based modelling, (ii) build a powerful mathematical tool to estimate fibrous biomaterials’ permeability tensor, (iii) obtain a deeper understanding on how microstructure affects transport property of fibrous biomaterials, and (iv) provide new tools for the noninvasive characterisation of fibrous biomaterials’ transport properties.

## 2. Materials and Methods

### 2.1. Geometric reconstruction

Brain WM is a typical anisotropic fibrous material; Fig. 1a & 1b show its transverse and longitudinal section views obtained by electron microscope [26, 27]. A key parameter to reconstruct the microstructure is the sizes of nerve fibres, which we have obtained experimentally by adopting the Focused Ion-Beam Scanning Electron Microscopy (FIB-SEM) technique (Fig. 1c) [28]. For the sake of data statistics and without loss of generality, the cross-sections of the nerve fibres were simplified to be circular (note ellipticity can also be captured and later introduced as a second order effect) and Fig. 1d presents the data of their diameters, which fits well to a log-normal distribution.

**Figure 1:**
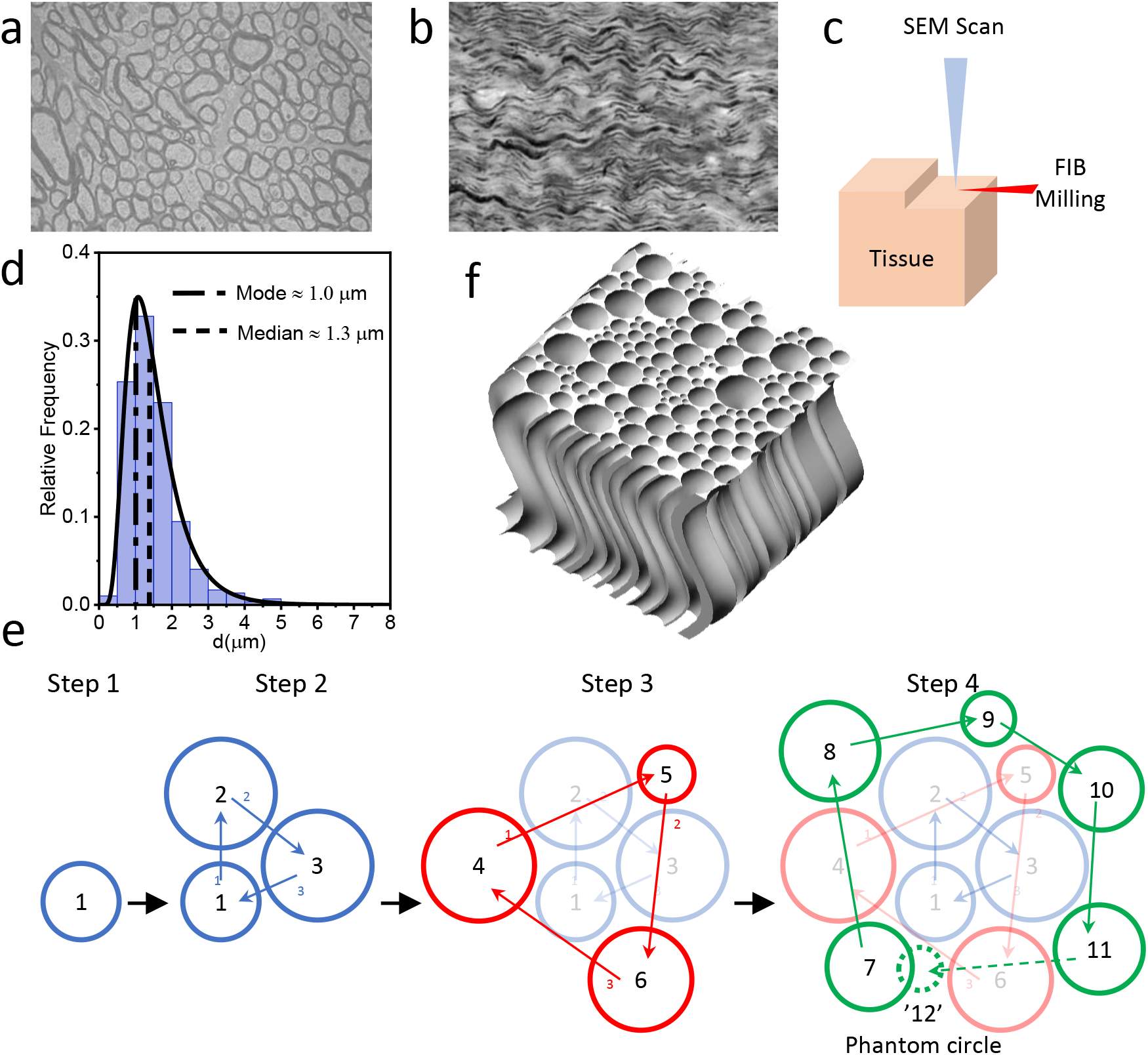
Strategies for microstructural reconstruction. **a**. Electron micrographs of the cross-sectional configuration of the brain white matter [26]. The disk-like structures are the cross-section of nerve fibres. **b**. Electron micrographs of the longitudinal configuration of the brain white matter, which shows the wavy shape of nerve fibres [27]. **c**. Schematic representation of Focused Ion-Beam Scanning Electron Microscopy (FIB-SEM) technique. Combining scanning and milling, we obtained the microstructural configuration of the ovine brain white matter every 150 nm, from which we measured the precise distribution of the nerve fibre’s diameter. **d**. Measured diameter distribution of ovine nerve fibres in the region of Corona Radiata. **e**. Schematic representation of the strategy to generate the randomly distributed cross-section of nerve fibres. **f**. A representative 3D geometry of ovine brain WM (Corona Radiata).

We then built an algorithm to randomly place these circles/disks in a rectangular domain to mimic the transverse section of the WM, which belongs to packing problems that have received extensive attention [29, 30, 31]. Three major challenges need to be overcome by the present algorithm: (i) the reported minimum porosity of brain tissue is 0.15 [32], which is close to the current limitation of documented algorithms for random disk packing [29]; (ii) diameters of all disks to be packed have been determined, so generating additional small disks to fill the pores left between large disks to reduce the porosity is invalid in this case; (iii) disk-disk and disk-boundary overlaps are not allowed. Under these conditions, it is challenging to randomly determine both the position and size of the disks while achieving a low porosity. We propose here a random-size-adaptive-position strategy to keep the randomness while acquiring low porosity. A schematic of the main strategy is shown in Fig. 1e, which contains five major steps:

- Step 1 & 2 - The core. We generate a random array of diameters based on data in Fig. 1d and number them in ascending order. Disk 1 is placed at the centre of a rectangular domain. Disk 2 is then placed on top of disk 1 within a specific distance. Note that we assume the tangent distance between any two disks is constant. Disk 3 is then placed to the right and tangential to disks 1&2.
- Step 3 - The outer layers. Outside the core, we start constructing the first outer layer. In each layer, we chain the outmost disks and place the new disks left and tangential to the chain in clockwise order. For example, for the first outer layer, we place the new disks tangential to disks in the core in a clockwise fashion from 1 to 2, 2 to 3 and 3 to 1.
- Step 4 - The 2D model. Step 3 is repeated until all the disks are placed. However, strictly following the chain may lead to disk-disk overlap, e.g. placing disk 12 left to 6 → 1. Under this condition, the algorithm will search for a smaller disk to fit to this position. If no smaller disk can fit, we will skip this position. In the case that disk and boundary overlap at the outermost layer, the same method is applied to solve the problem.
- Step 5 - The 3D model. Step 4 generates a large cross-sectional domain, but we may just use a part of it to run simulations, which depends on the size of the Representative Volume Element (RVE), which will be discussed in Section 2.3, so we cut a rectangular domain and extrude it along a wavy line to generate the 3D geometry, as shown in Fig. 1f. The wavy line is defined by the average length and tortuosity (*τ*: the ratio of the arc length to the distance between the two endpoints) of the fibre, which, for specific brain axonal structures, are measured as 15 μm and 1.1, respectively [28].

### 2.2. Mathematical expressions for the ϕ – δ – κ relationships

In many studies, the porosity-permeability relationship (*κ* = *f* (*ϕ*)) can be analytically obtained via solving the Navier-Stokes (NS) equations in an idealised and periodic geometry, as exemplified by Fig. 2a [33, 14]. In biomaterials, however, the randomness of the fibre size and position makes it impossible to adopt analytical methods. Inspired by the scaling rule by Clague [34], we write the permeability as a function of the half distance between the disks (i.e. *κ* = *f* (*δ*), the parameter *δ* is shown in Fig. 2). Under the low Reynolds number regime, which holds for the majority of biomaterial and biological tissues [35], Clague performed a simple analysis of the Stokes flow equations:

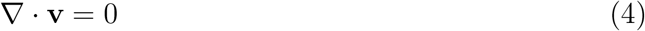

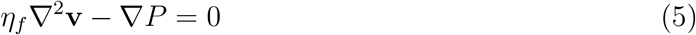

and deduced the following scaling estimate for Stokes flow in the direction perpendicular to the fibres, as shown in Fig. 2b:

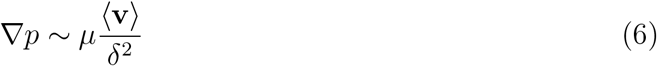

or

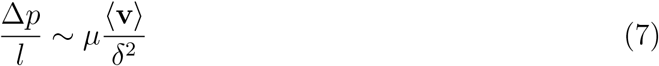

where **v** is the flow velocity, *η_f_* is the fluid dynamic viscosity (0.8 mPa·s in this study [36]), *p* is the hydraulic pressure, 〈**v**〉 and *l* are the characteristic velocity and length, respectively. Substituting Eq. 7 to Darcy’s Law:

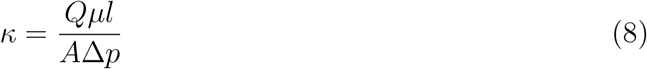

where *Q* is the flow rate and *A* is the cross-sectional area of the domain, we can write:

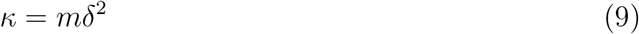

where *m* is a coefficient dependent on the pore-scale microstructure.

**Figure 2:**
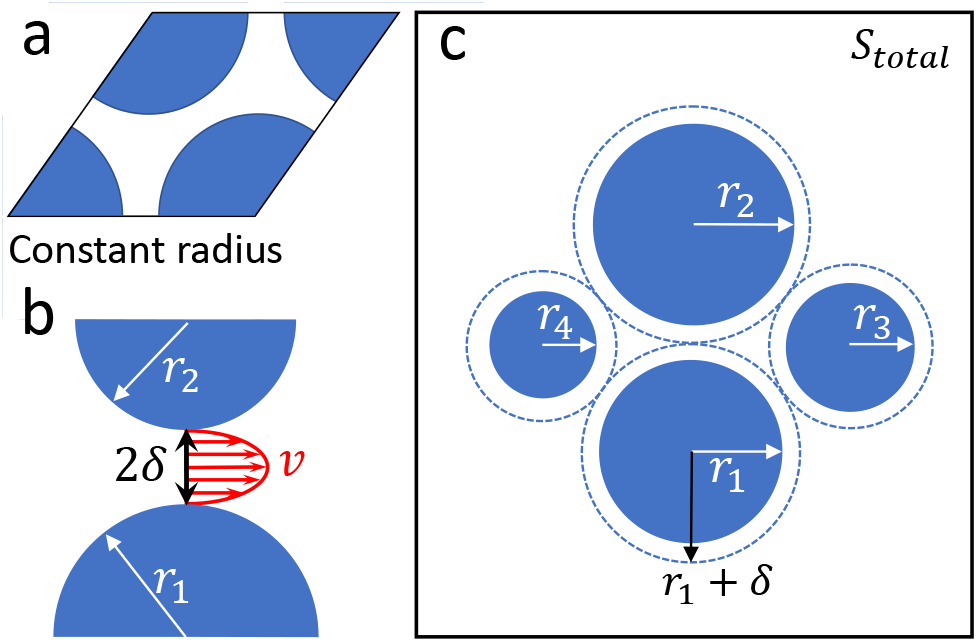
**a**. An example of idealised geometry for porosity-permeability relationship studies. **b**. Schematic representation of mathematically deriving the *δ* – ***κ*** relationship. **c**. Schematic representation of mathematically deriving the *δ* – *ϕ* relationship.

In line with what has been suggested by other researchers [37, 38], we treated the WM as transversely isotropic; this implies that the nerve fibres and ISF behaviours on the plane perpendicular to the main fibre trajectory are directions independent. We can thus define the permeability tensor as ***κ*** = diag(*κ*_⊥_, *κ*_⊥_, *κ*_∥_) and the perpendicular component is:

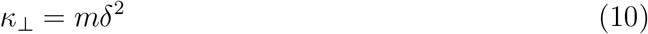

which shows that when the fluid flows perpendicular to the nerve fibres, the permeability is proportional to the squared half distance between the fibres; and when there is no gap between the fibres (i.e. *δ* = 0), the permeability reduces to zero as no flow can develop between the fibres, which provides a good representation of the behaviour expected using physics-based arguments. When the fluid flows parallel to the nerve fibres, however, the permeability will not fall to zero even if *δ* = 0, because the void spaces between the fibres can be the flow pathway. Therefore, the format of *δ* – *κ* relationship needs to be:

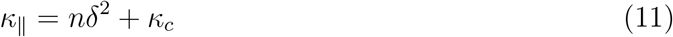

where *n* is similar to *m*, a coefficient dependent on the pore-scale microstructure, and *κ_c_* is the critical parallel permeability when *δ* = 0.

To write *κ* as a function of porosity (*ϕ*), we then need to find the relationship between *δ* and *ϕ*. Let’s define *S_total_* as the area of the whole domain, *S_c_* as the critical void area when *δ* = 0, *r_i_* as the radius of each fibre and *N* the number of fibres, as shown in Fig. 2c. We then have:

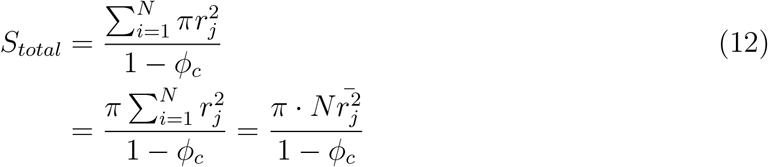

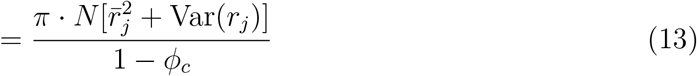

where 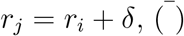 is the mean value, and Var(·) is the variance:

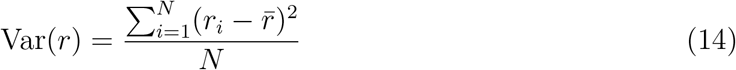

The porosity can then be written as:

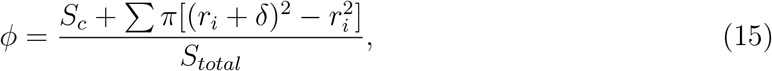

which rearranges to give

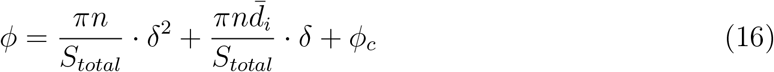

where *ϕ_c_* = *S_c_/S_total_* and *d_i_* represents the fibre diameters. If we write Eq. 16 in the form

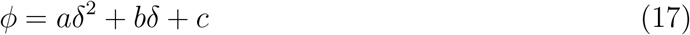

we then have

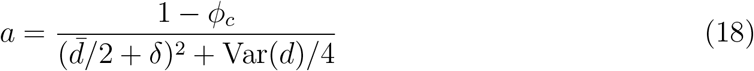

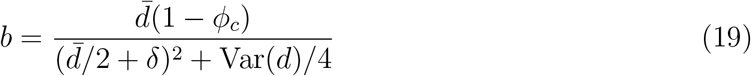

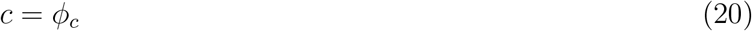

By combining Eqs. 10, 11, 17, the relationship between *ϕ* and *κ* can be obtained, but we still need to know the coefficients *m, n*, and the values of *a, b, c* (i.e. 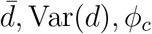. In this study, 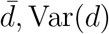 can be easily obtained from Fig. 1d, so we are only left with *m, n,ϕ_c_* unknown. *ϕ_c_* can be obtained by running the algorithm developed in Section 2.1 to reconstruct transverse sections with *δ* = 0. Since *m* and *n* link the geometric parameter to the hydraulic permeability, they can be characterised by running computational fluid dynamics (CFD) simulations in the reconstructed 3D geometries.

### 2.3. CFD and RVE

According to Darcy’s law (Eq. 8), the value of *κ* can be calculated from the flow rate (Q) and pressure drop (Δ*p*) in a given domain. As these two values can be obtained by solving Stokes equations (Eqs. 4, 5) in the domain, we run computational fluid dynamics (CFD) in the representative volume elements (RVEs) with different *δ* to characterise m and n. In the cases of parallel flow (see Fig. 3a), the top surface of the RVE is set as the inlet boundary with the inlet pressure of 10 Pa, the bottom surface is set as the outlet boundary with the outlet pressure of 0 Pa. Note that *κ* is independent of the pressure drop, so the inlet pressure and outlet velocity can be arbitrarily chosen, i.e. the calculated *κ* will not change if the inlet pressure and outlet pressure are set to be any other values [39]. The boundaries of nerve fibres are set as no-slip. In the cases of perpendicular flow (see Fig. 3b), the right surface of the RVE is set as the inlet boundary with the inlet pressure of 10 Pa, the left surface is set as the outlet boundary with the outlet pressure of 0 Pa.

**Figure 3:**
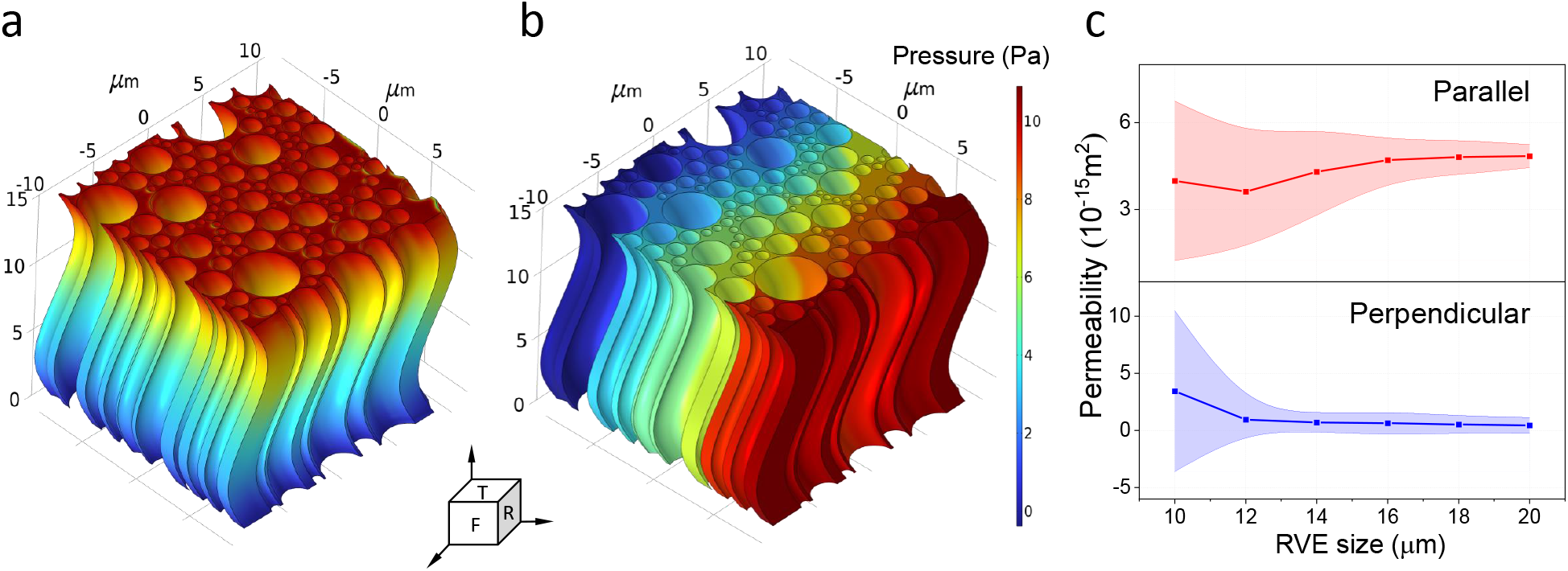
CFD setup and RVE size. **a**. A representative result of pressure distribution in the RVE when the flow is parallel to the nerve fibres. The top (T) and bottom surfaces are the inlet and outlet boundaries, respectively. **b**. A representative result of pressure distribution in the RVE when the flow is perpendicular to the nerve fibres. The right (R) and left surfaces are the inlet and outlet boundaries, respectively. **c**. The relationship between RVE size and calculated permeability in two directions. The shadow areas are the standard derivations (SD) of the results.

The size of RVE (side length of the top and bottom surfaces in this study) has a significant effect on simulation results, as shown in Fig. 3c. When the RVE size is relatively small, the calculated values of permeability using different RVEs cannot converge to a stable value, but the instability decreases with the RVE size. Based on the results in Fig. 3c, we choose 18 μm as the RVE size. It is worth mentioning that the RVE sizes we obtained here are very similar to those obtained by [40].

### 2.4. Experimental data and Poroelastic modelling

We verified the newly proposed ***κ*** = *f* (*ϕ*) formula by a group of brain tissue infusion experiments, where Phosphate-buffered saline (PBS) was injected into the brain WM tissues both along and perpendicular to the nerve fibres, as shown in Fig. 4a, and the corresponding changes of permeability tensor were measured [41]. The box charts in Fig. 6 show the experimental results. We built an anisotropic poroelastic model and integrate the ***κ*** = *f* (*ϕ*) formula to simulate the infusion experiments.

**Figure 4:**
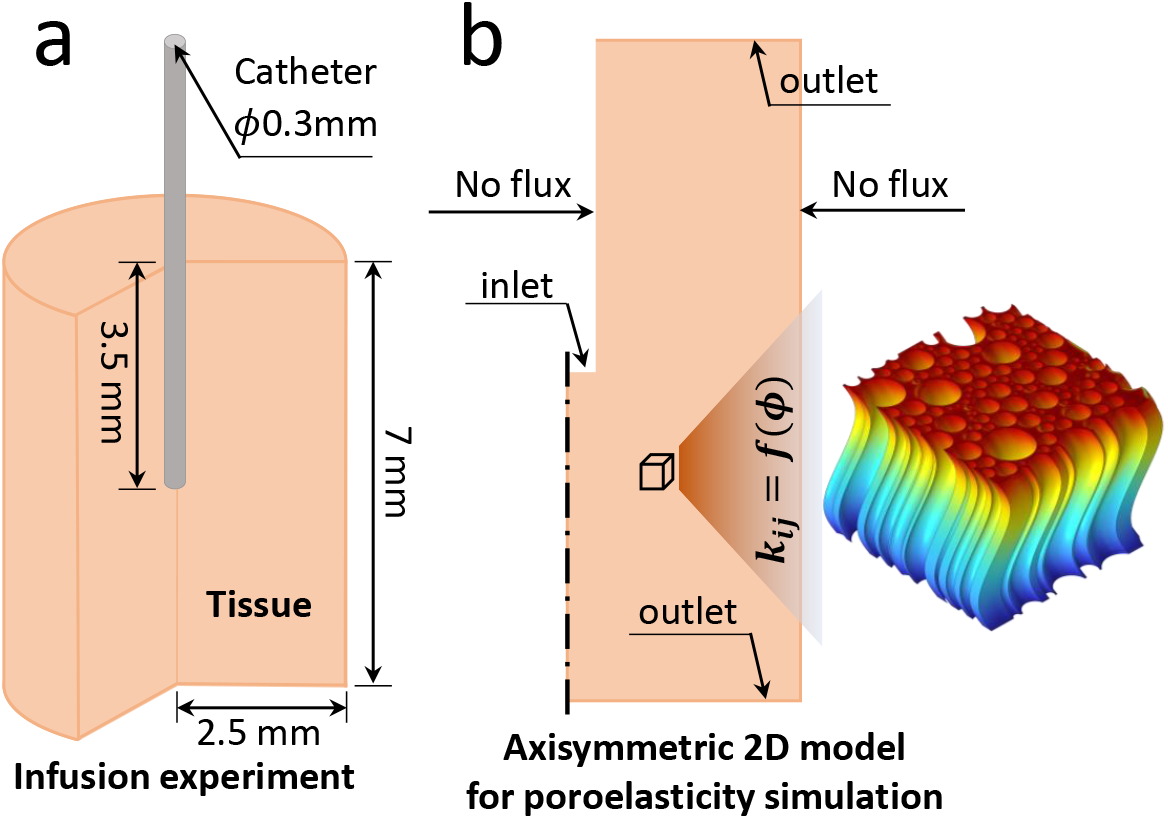
**a**. Schematic representation of the brain tissue infusion experiment. **b**. Schematic representation of the Geometric model, boundary conditions, and the method of integrating the ***κ*** = *f* (*ϕ*) formula for the anisotropic poroelastic model.

**Figure 5:**
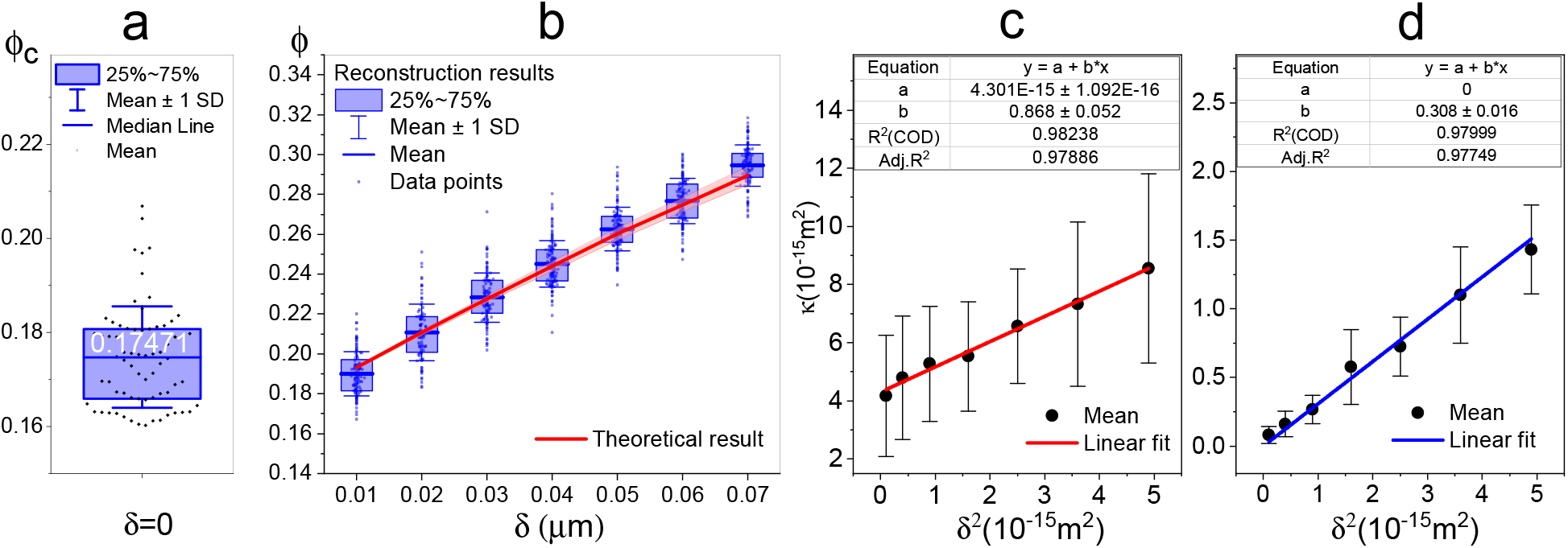
*δ* – *ϕ* – *κ* relationships. **a**. Critical porosity (*ϕ_c_*) of the fibrous porous medium when *δ* = 0. **b**. *δ* – *ϕ* relationships obtained by direct measurement of the reconstructed RVEs (box chart) and using the newly developed theory (shadowed line). The central line shows the mean values while the shadow denotes the SD. **c**. Characterisation results of *δ*^2^ – *κ*_∥_ relationship of parallel flow. the error bars denote SD of the results in each group. The table on the top shows the characterised formula. **d** Characterisation results of *δ*^2^ – *κ*_⊥_ relationship of perpendicular flow.

**Figure 6:**
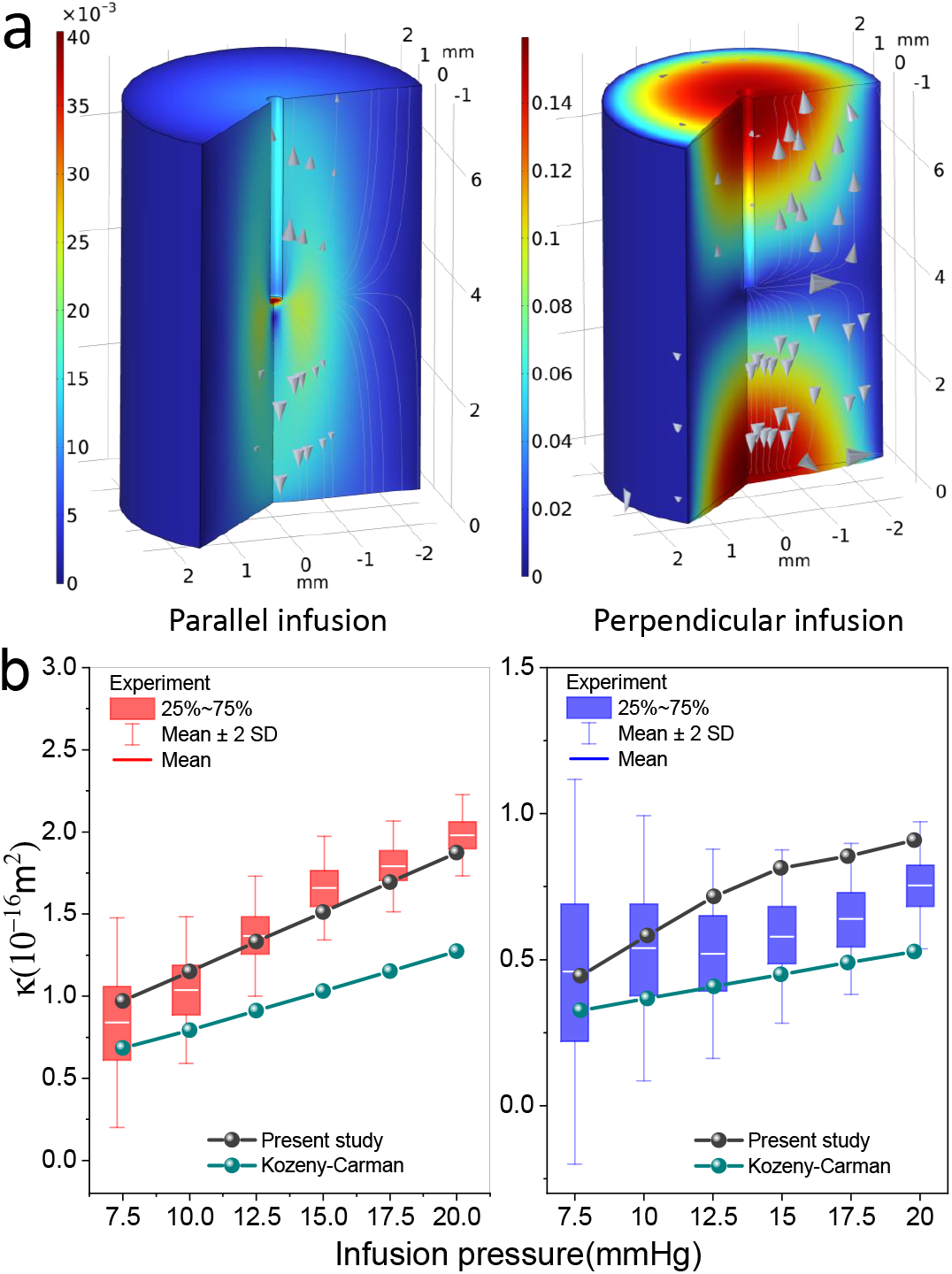
Comparison between experimental results and theoretical results. **a**. Poroelastic modelling result of brain tissue infusion. The contours show the tissue deformation (mm) while the vectors denote the flow direction. **b**. Infusion pressure-permeability tensor relationship. The box charts are experimental results, while the lines present the theoretical results. In both figures, ***left*** is for parallel infusion and ***right*** is for perpendicular infusion.

In the poroelastic theory, the porous domain is divided into the solid phase and the fluid phase. As brain WM is composed of nerve fibres, interstitial fluid (ISF), and extracellular matrix (mainly lecticans and proteoglycans) [42], we assume the nerve fibres as the solid phase and the rest as the fluid domain. We can then write the stress relationship as:

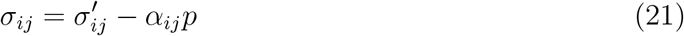

where *σ_ij_* is the total (tissue) Cauchy stress, 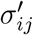 is the Cauchy stress of nerve fibres, *α_ij_* is the Biot-Willis coefficient tensor, and *p* is the interstitial hydraulic pressure. To more precisely capture the fibre’s compressive behaviour, We used a realistic hyper-elastic model (Eq. 22) to describe the nerve fibres and the fibre stress can be written as Eq. 23.

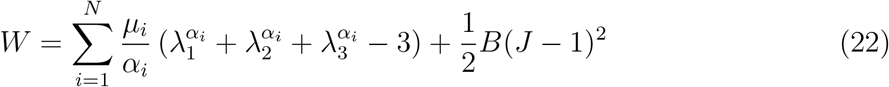

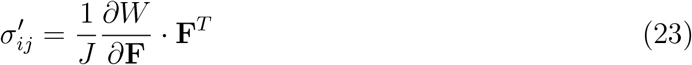

where *W* is the strain-energy density function that is expressed in terms of the principal stretches *λ_i_,i* = 1, 2, 3; *N, μ_i_, α_i_* are material constants with the values of 1, 281.84 Pa, and 6.33, respectively [27]; *B* is the bulk modulus, and 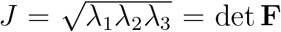 is the volumetric deformation, and **F** = *f_ij_* = *∂***x***_i_*/*∂***_X_***_j_* is the deformation gradient tensor. The updated porosity due to nerve fibres’ compression can be then written as:

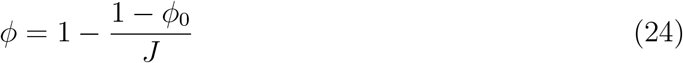

where *ϕ*_0_ is the initial porosity without fibres’ compression (~ 0.2 for brain tissues [32]). Integrating the updated porosity and permeability tensor to Darcy’s Law, we can then solve the flow velocity:

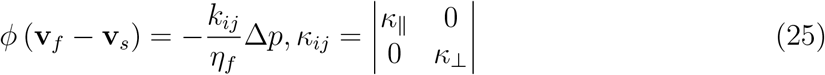

where **v***_f_* is the flow velocity, **v***_s_* is the deformation velocity of the solid phase.

## 3. Results

### 3.1. A versatile microstructure generator for fibrous material: MicroFiM

Based on the algorithm presented in Section 2.1, we built a software application with MATLAB, which is able to generate closely and randomly packed porous domains. By inputting the distribution of fibre diameters (currently supports Lognormal Distribution, but the source code can be easily changed to support any other distributions), the approximate distance between fibres and the size of the domain, the software will automatically generate the geometric information of the porous media. The App and source code are available in the Supplementary material and the manual is available in Appendix A.

Prevailing opinions suggest that the distance between neurons ranges from ~30 nm to ~100 nm [43, 44]; the extracellular space (ECS), accordingly, occupies normally 18% ~ 30% of brain tissues [32]. The parameter values of the generated microstructures, underpinned by our measurement of neuron fibre diameter distribution (Fig. 1d), coincide exactly with the ranges. First, by reconstructing 70 RVEs with *δ* = 0 and measuring their porosity, we obtained ϕ_c_ ≈ 0.175 (Fig. 5a), which agrees that the lowest ECS volume fraction should be higher than 0.18. Second, we reconstructed 7 groups of RVEs (each group contains ~100 RVEs) with the fibre distance in the range of 20 nm ~ 140 nm. The porosity of these RVEs is, accordingly, between 0.17 ~ 0.32, as shown by the box chart in Fig. 5b.

### 3.2. δ – ϕ relationship

The half-distance between fibres (*δ*) bridges the tissue porosity (*ϕ*) and permeability tensor (***κ***) in the present study, so we first characterised the formulas of *ϕ* = *aδ*^2^ + *bδ* + *ϕ_c_* and ***κ*** = diag(*mδ*^2^, *nδ*^2^ + *κ_c_*), i.e. the coefficients of *ϕ_c_, m, n, κ_c_*, according to Section 2.2.

Substituting the mean value 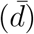 and variance (Var(*d*)) of nerve fibre diameters of each RVE and the *ϕ_c_*, which has been available in Section 3.1, to Eqs 19, 20, we obtained the theoretical results of the *δ* – *ϕ* relationship of each RVE presented by the shadowed red line in Fig. 5b. The central line shows the mean value and the shadow shows the standard deviation (SD). Comparison between the measured results and the theoretical result demonstrates the validity of Eqs. 16-20 and the precision of *ϕ_c_*. It is worth mentioning that compared with *ϕ_c_*, 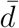 and Var(*d*) of fibre diameters are more easily available by imaging techniques, so we characterised *ϕ_c_* in this study which can be directly adopted for brain tissue. For other fibrous materials, given the diameter distribution, the App can easily characterise their specific *ϕ_c_*.

### 3.3. δ – κ relationship

We then run CFD in these RVEs and calculated their individual permeability in both directions based on Darcy’s law. The results of parallel flow and perpendicular flow are shown in Fig. 5c and d, respectively, with error bars. It shows that in both directions, the mean value of *κ* is proportional to *δ*^2^, which matches our theory expressed by Eqs. 10 and 11. By linearly fitting the mean values, we finally characterised *m, n, κ_c_* and obtained the following formulas:

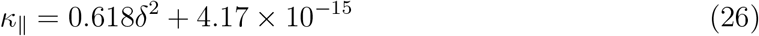

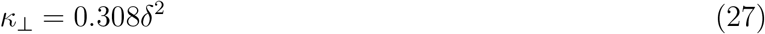

### 3.4. Application and validation

Fig. 6a shows the results of modelling the brain tissue infusion experiments with the anisotropic poroelastic model built in Section 2.4 underpinned by the newly developed ***κ*** = *f* (*ϕ*). While the streamlines show the flow direction, the contours show the tissue deformation. Due to the existence of a plastic tube wrapping and supporting the tissue, the fluid cannot flow out via the outer boundaries of the tissue, so the streamlines turn to the axial direction when reaching the outer boundary of the tissue. In addition, streamlines in the right figure (infusion is perpendicular to nerve fibres) turn more sharply to the axial direction compared with the situation in the left figure (infusion is parallel to nerve fibres), this is because the flow can be more easily developed along the nerve fibres. When applying local hydraulic pressure, nerve fibres tend to deform laterally because the fluid can flow freely along nerve fibres. Consequently, as shown in the contours, the parallel infusion does not introduce obvious tissue deformation while the perpendicular infusion extrudes the tissue.

In Fig. 6b, we compared the simulation results (lines with symbols) against the experimental results (boxes). It shows that adopting the Kozeny-Carman formula seems to be able to provide a good prediction under relatively low infusion pressure but leads to significant error under high pressure, especially in the scenario of parallel infusion. The lack of consideration of the flow along fibres should be the culprit. By contrast, the newly developed formula in the present study can better predict the infusion in both directions, but the result in perpendicular infusion is slightly higher than the mean values of experimental measurements under high pressure. The reason is that in the situation of perpendicular infusion, apart from being compressed, the nerve fibre will also undergo lateral deformation. While compression of the solid phase provides more space for the fluid flow thus increasing permeability, the lateral deformation of nerve fibres may drive the nerve fibres to approach each other, close the flow pathway and decrease the permeability. Given that the formulas of **κ** = *f* (*ϕ*) can only consider the porosity change caused by compression of the solid phase, the predicted permeability under perpendicular infusion should be higher than the real situation. To prove this hypothesis, we added spring foundations to the outlet boundaries (see Fig. 4b) to reduce the lateral deformation and mimic the situation of closing gaps between nerve fibres. See more details in Appendix B.

### 3.5. Effects of fibrous microstructure on the ϕ – κ relationship

To obtain a clear understanding of how the fibrous microstructure and the randomness of the fibre size affect the porosity-permeability tensor relationship, we further conducted parametric studies on the fibre length, tortuosity and diameter distribution to explore their effects.

#### Fibre length and tortuosity

Changes in fibre length and tortuosity do not change the cross-sectional configuration of the material, so only changes in parallel permeability need to be considered in these cases. We generated 7 new groups of RVEs with different lengths and tortuosities, see Fig. 7a and 7b for the specific values. Results show that fibre length alone has a very limited effect on the porosity-permeability relationship. We then chose one cross-sectional configuration with different length, and found that although the permeability values decrease slightly with the length, the deviations are within 3% (see Table 1).

**Figure 7:**
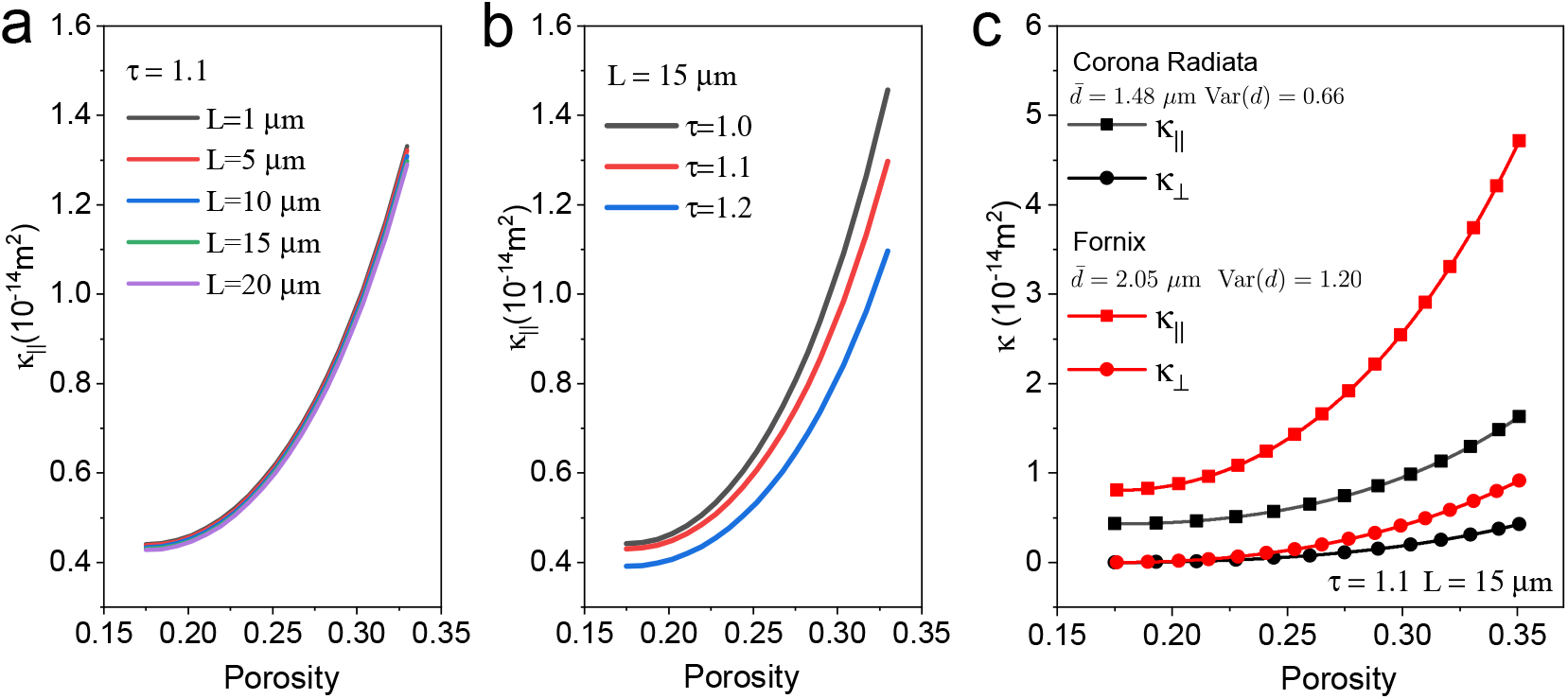
Microstructural effects on the porosity-permeability relationship. **a**. Effect of fibre length on the *ϕ* – ***κ*** relationship. Fibre tortuosity is 1.1. **b**. Effect of fibre tortuosity on the *ϕ* – ***κ*** relationship. Fibre length is 15 μm. **c**. Effect of fibre diameters’ distribution (region) on the *ϕ* – ***κ*** relationship.

**Table 1:**
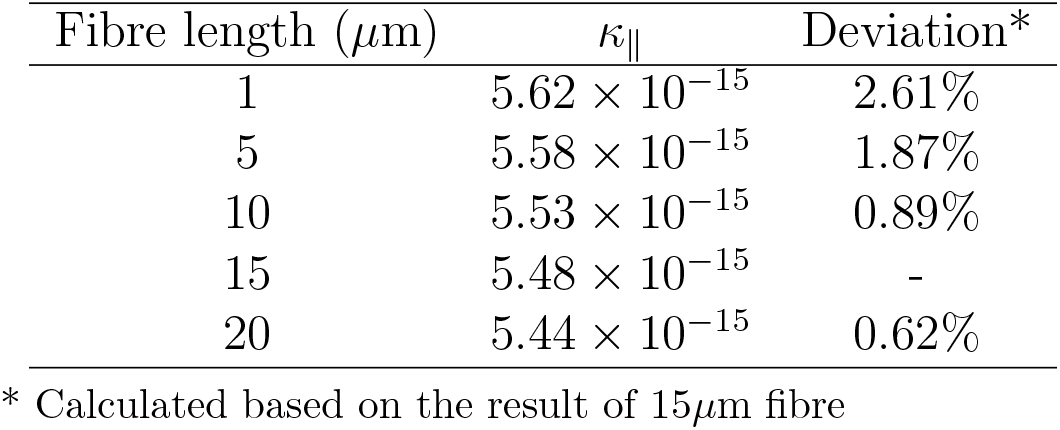
Relationship between fibre length and parallel permeability

On the contrary, the effect of tortuosity is considerable. Fig. 7b shows that a larger tortuosity leads to a smaller parallel permeability and also a slightly slower increase of parallel permeability with porosity; these effects also increase with the tortuosity. This suggests that for highly curved fibres, we should also incorporate tortuosity into the formula of the porosity-permeability tensor relationship. This will be explored in more detail in future work.

#### Fibre diameter distribution

Different from the above situations, changes in fibres’ diameter distribution also change the cross-sectional configuration of the material, so we studied both components of the permeability tensor here. In this case, we chose another area of ovine brain white matter (Fornix) and adopted the same protocol as dealing with the region of Corona Radiata (Fig. 1). Appendix C, which shows the processing details of Fornix, again demonstrates the validity of applying the theory established in Section 2.2 to Fornix. Comparisons of the porosity-permeability tensor relationship between Corona Radiata and Fornix show that with larger mean value and variance of fibre diameters, Fornix has larger permeability in both directions than Corona Radiata under the same porosity, and the difference increases with the porosity. By comparing Fig. 5b and S2c, we further found that both theory and direct measurement show that with the same distance between fibres, Fornix has a smaller porosity than Corona Radiata. These two findings show that while we may intuitively acknowledge that a higher porosity means higher permeability, the real situation is more complex and may be counter-intuitive, which further underscores the importance of defining a reliable constitutive model for the porosity-permeability relationship of the fibrous domain formed by fibres with random size and position.

## 4. Discussion

While extensive studies have shown the significant impact of the microstructure on mass and fluid transport processes in biomaterials [45, 46], our recent experimental and imaging results have further emphasised this relationship and raised the more complicated anisotropic issue existing more widely in biomaterials and biological tissues e.g. brain white matter [41, 47]. Although we have successfully unravelled the mechanisms behind the complex interactions between the microstructure and transport processes via theoretical modelling [6, 48], the practical utilisation of this relationship to gain an improved understanding of the functions of biological tissue and design capability of advanced biomaterials is still difficult due to the lack of an explicit definition of the relationship between the microstructure and transport properties. The framework developed in this study provides a solution to bridge this gap, specifically between the microstructure and the anisotropic transport property of fibrous biomaterials as well as other types of composite materials.

Mechanical behaviours are important for biological tissues and biomaterials [49]; their characterisation usually rely on precise definitions of mechanical and spatial interactions between the solid phase and fluid phase. Consequently, the poroelastic theory is widely adopted in modelling the mechanical behaviours of biological tissues and biomaterials and the precision of modelling results relies on the definition of the porosity-permeability relationship. Due to the lack of a porosity-permeability tensor relationship that can explicitly consider the microstructural complexity of fibrous biological tissue and biomaterials, modelling their anisotropic mechanical behaviours is still an outstanding problem. Although the scope of this study provides only the porosity-permeability tensor relationship of brain white matter, we can adapt the theory, tools, and protocols established here to build the porosity-permeability tensor relationship of other similar fibrous biomaterials and therefore pave the way for a more precise prediction of their mechanical properties.

Extensive studies in the recent decade have shown the direct links between the fluid & mass transport in the brain and brain diseases, such as Parkinson’s, Alzheimer’s and stroke; particularly since the dysfunctions of these transport processes can reduce the waste clearance and spread the toxic components or pathogens [5]. Whereas how the complex brain microstructure guides the fluid flow and how the unhealthy microstructure leads to the abnormal fluid flow are largely unknown and unpredictable. Aided by advanced medical imaging techniques, e.g. neurite orientation dispersion and density imaging (NODDI) [50], which can calibrate the distribution of porosity in the brain, the newly developed quantitative relationship between porosity and permeability tensor of brain white matter will be able to draw the map of whole brain fluid transport. And therefore potentially helping to detect brain disorders by monitoring the brain flow status. The potential success of the emerging drug delivery techniques, which deliver drugs directly into the brain tissue by infusion, also depends on the understanding of transport properties of the brain [13]. Successful application of this newly proposed mathematical formula and supporting methods will be able to play important role in planning the drug delivery route in the brain. One potential limitation is that these applications need to be underpinned by a database of nerve fibre diameters’ distribution in different regions of the human brain as the results in Fig. 7c suggest significant differences of permeability tensor exist in the different brain region. Fortunately, the protocol to build this database has been established in Ref. [28].

## 5. Conclusions

In the present study, we have (i) built a geometry generator for closely and randomly packaged fibrous domains and can reach a low porosity down to 0.16. By generating the complex and semi-realistic microstructure, this tool will enable the community to carry out various types of modelling to capture biomaterial behaviours and gain a deeper understanding of microstructure-function relationships; (ii) built a validated mathematical theory and comprehensive modelling protocol to link the microstructural information of fibrous biomaterials and biological tissues to their anisotropic permeability tensor, which fills the gap of lacking a precise porosity-permeability tensor relationship to accelerate biomaterial design and non-invasive characterisation; (iii) proposed a validated mathematical formula of the porosity-permeability tensor relationship of brain white matter, which will help to precisely model fluid transport in the brain tissue and plan the drug delivery route in the brain.

## Acknowledgements

This project has received funding from the European Unions Horizon 2020 research and innovation programme under Grant Agreement No. 688279. Daniele Dini would like to acknowledge the support received from the EPSRC under the Established Career Fellowship Grant No. EP/N025954/1. Li Shen thanks the Engineering and Physical Sciences Research Council (EPSRC) for a Postdoctoral Fellowship (EP/V005073/1).

