## Supplementary Information for "Porosity-permeability tensor relationship of closely and randomly packed fibrous biomaterials and biological tissues: Application to the brain white matter"

#### Appendix A: Manual of MicroFiM

MicroFiM is a MATLAB application build based on MATLAB R2020b [1]; Fig. S1 shows the GUI. To meet the requirement of the present study, the current version supports generating fibres with their diameters following lognormal distribution. For other types of distributions, the users could modify the source code. The mode value, median value, and mean value of the fibre diameter are needed to be input. The nodes 'No. of Geometry' and 'No. starts from' are designed for generating multiple geometries with designated labels. 'RVE size' and 'Fibre Distance' can be freely chosen based on the users' requirements. The generated microstructural information is stored as .xlsx files for general use. The users can choose the path to store the files in 'Result Directory'. Before running the application, the users are suggested to click 'Check' to confirm if the input information is correct. All the processing information will be updated in the 'Status' window.

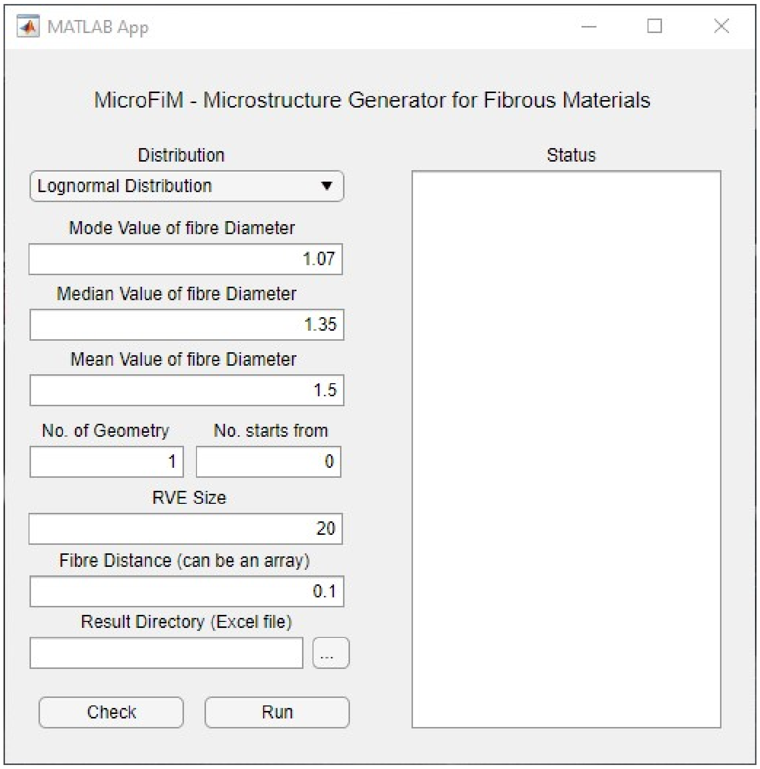


Figure 1: Graphical User Interface (GUI) of the geometry generator.

### Appendix B: Poroelastic simulation results under different boundary conditions

The black lines in Fig. S2 show the results of modified boundary condition. It shows that modifying this boundary condition does not affect the result in parallel infusion, because the deformation on outlet boundaries is very limited. In the perpendicular infusion situation, this modification reduces permeability when infusion pressure is high, because a higher pressure leads to larger lateral deformation and narrower flow pathway thus more significantly reduce the perpendicular permeability.

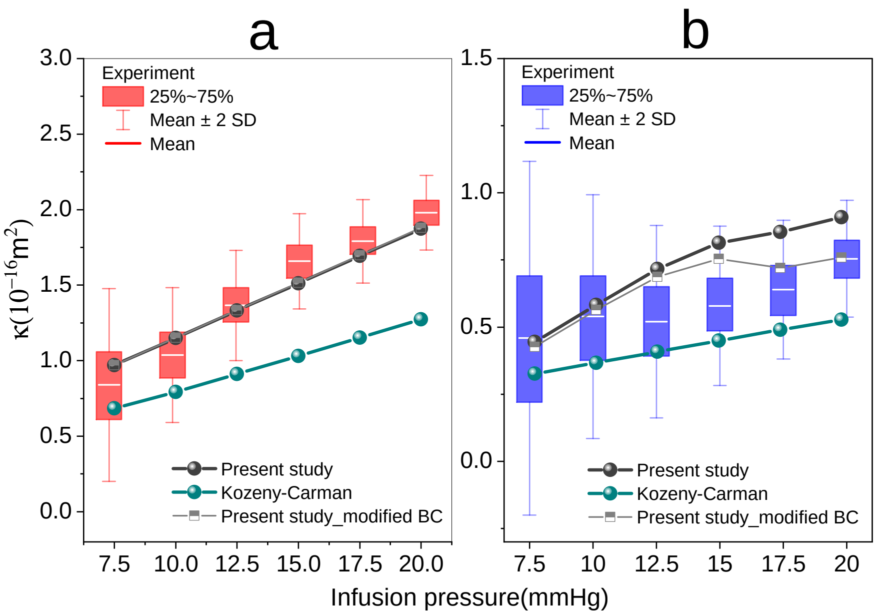


Figure 2: Infusion pressure-permeability tensor relationship. The box charts are experimental results, while the lines present the theoretical results. **a**. Parallel infusion and **b**. Perpendicular infusion.

### Appendix C: Data processing for Fornix

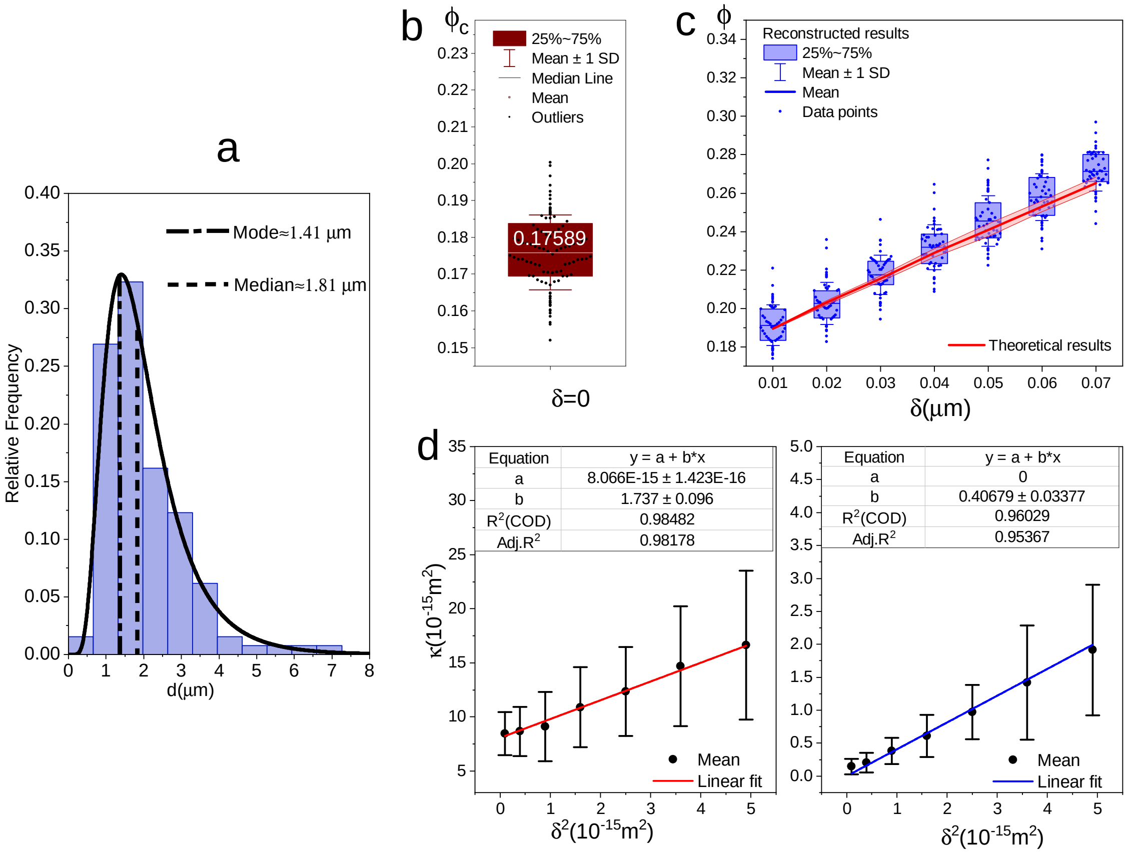


Figure 3: Geometric information and data fitting of Fornix. **a**. Diameter distribution of nerve fibres in Fornix. **b**. Characterisation of $\phi_C$ for Fornix. **c**. The $\delta - \phi$ relationship of Fornix. **d**. The $\delta^2 - \kappa$ relationship in parallel direction. **e**. The $\delta^2 - \kappa$ relationship in perpendicular direction.

#### References

[1] MATLAB, version R2020b, The MathWorks Inc., 2020.
